# Biology of healthy aging: Biological hallmarks of stress resistance-related and unrelated to longevity in humans

**DOI:** 10.1101/2024.07.19.604380

**Authors:** Komalpreet Badial, Patricia Lacayo, Shin Murakami

## Abstract

Stress resistance is tightly associated with longer and healthier lifespans in various model organisms, including nematodes, fruit flies, and mice. However, we lack a complete understanding of stress resistance in humans, and therefore, we investigated how stress resistance and longevity are interlinked in humans. Using more than 180 databases, we identified 541 human genes associated with stress resistance. The curated gene set is highly enriched with genes involved in cellular response to stress. The Reactome analysis identified 398 biological pathways, narrowed down to 172 pathways, using a medium threshold (p-value < 1 × 10^−04^). We further summarized into 14 pathway categories, e.g., cellular response to stimuli/stress, DNA repair, gene expression, and immune system. There were overlapping categories between stress resistance and longevity, including gene expression, signal transduction, immune system, and cellular responses to stimuli/stress. The categories include the PIP3-AKT-FOXO and mTOR pathways, known to specify lifespans in the model systems. They also include the accelerated aging syndrome genes (WRN and HGPS/LMNA), while the genes were also involved in non-overlapped categories. Notably, nuclear pore proteins are enriched among the stress resistance pathways and overlap with diverse metabolic pathways. This study suggests that stress resistance is closely linked with longevity pathways, but not entirely identical. While most longevity categories intersect with stress-resistance categories, some do not, particularly those related to cell proliferation and beta-cell development. We also note inconsistencies in pathway terminologies with aging hallmarks reported previously and propose them to be more unified and integral.

## 1. Introduction

Aging is a complex biological process characterized by gradual changes, mostly declines in biological function and increased vulnerability to external and internal stress over time. The age-related changes and modifications occur in macromolecules, leading to genetic and epigenetic changes at multidimensional levels, ranging from molecules, cells, extracellular regions, tissues, and the whole body. The major challenge in the field of aging has been to identify the causes of aging and distinguish their consequences. Herman (1956) focused on oxidative stress, such as reactive oxygen species, namely ROS (OH and HO_2_), as a possible cause of aging (the free radical theory of aging) [1]. Mutations in ROS scavenging enzymes, such as superoxide dismutase genes, however, have little or modest effects on lifespans [2–5]. With a wide variety of ROS functions, it has been experimentally difficult to prove that oxidative stress is the sole cause of aging despite its deleterious nature as single stress, and mounting evidence suggests that oxidative stress may play more specific roles in age-related diseases [6–7]; Similarly, thermotolerance dependent on HSF-1 can be separated from life extension in worms [8]. Importantly, genetic and pharmacological screenings for single stress resistance (resistance to a single form of stress) have yielded those that confer life extension and those that do not confer life extension [9–11], suggesting two types of stress resistance, longevity-related and -unrelated. Thus, considering single stress as a sole cause of aging has limitations.

In the model systems, including worms and mice, life-extending mutants show increased resistance to various stressors, including oxidative stress, heat, ultraviolet light, and DNA-damaging agents, among others [Reviewed in Ref.12-15]. This resistance to multiple forms of stress (multiplex stress response and resistance, or multiplex stress resistance, in short) is better associated with life extension. The original concept of multiplex stress resistance was first introduced, reading “The finding that all Age (life-extending) mutations show resistance to multiple distinct stresses suggests the possibility that a common molecular mechanism may regulate the response to all three stressors (oxidative stress, heat and UV) [16].” The term, multiplex stress resistance, was later defined when the finding was extended to the cells of long-lived dwarf mice [17–18]. Furthermore, multiplex stress resistance is reduced during aging in worms [19] and the pathways for stress responses are needed for life extension [13,15] (Murakami, 2006; Soo et al., 2023). However, stress resistance does not explain all aspects of aging [20–21], which suggests the transitional phase before advanced aging [22–23]. Taken together, we still do not have a clear picture of what biological pathways contribute to stress resistance and how they are related to longevity.

Here, we investigated the biological pathways related to stress resistance in humans. We extracted a list of genes associated with stress resistance in humans and investigated biological pathways with a medium confidence level (p-value < 1 × 10^−04^). Our findings revealed that pathways associated with stress resistance overlapped most of the longevity pathways previously identified [24]. However, the remaining categories did not overlap with longevity.

## 2. Materials and Methods

### Nomenclatures

We define “single stress resistance” that refers to the ability of an organism to withstand a specific type of stress, such as oxidative stress, heat, or chemical exposure and “multiplex stress resistance”, on the other hand, refers to the ability of an organism to withstand multiple forms of stress. To avoid complex ontology nomenclatures, we used the terms as previously described [25–26]. Briefly, the Reactome ontology pathways are composed of two or more levels, ranging from the most general pathways (referred to as pathway categories or simply, categories) divided into more specific pathways to specific pathways (referred to as specific pathways or simply pathways) among others. For example, a pathway category of autophagy has in the order of autophagy (pathway category) > macroautophagy (more specific pathway) > selective autophagy (specific pathway). We have grouped all ontology pathways into each general pathway, namely pathway categories.

### Datasets

A pool of the curated gene set was initially identified using Genecards.org; it is a bioinformatic tool that ciphers through more than 180 scientific databases by inputting a search query and providing a list associated with human genes [26] (last accessed on 3/17/2023). The genes relevant to stress resistance were searched using the platform, Elastic search 7.11 (https://www.genecards.org/Guide/Search#relevance) (last accessed on 3/17/2023), which is based on the Boolean model to retrieve matching documents and the relevance score calculated by Lucene’s practical scoring function. The base keywords, “stress resistance” AND “humans” were used to search the genes with the relevance score. The relevance score did not come with confidence levels, and thus we further validated the relevance to stress resistance, using Reactome and STRING-DB as described in the text and also described previously [24].

### Gene Ontology

The Reactome analysis and STRING-DB analysis were performed as described previously [25]. The Cytoscape plugin Reactome FIViz classified and assembled the exported gene list into biological pathways. Reactome utilizes a systematic accumulation of pathway databases that performs analyses of biological pathways. The biological pathways are organized in a stratified fashion, often with sub-pathways. Sub-pathways are strings linked to an arching pathway, therefore hit genes can overlap with different arching pathways. The exported gene list associated with stress resistance was inputted into the Reactome FIViz program. The program then executed and carried out a range of gene set analyses at a False Discovery Rate (FDR) of 0.05. The research team independently conducted the task to cross-reference and validate the retrieved gene set analysis. We used more stringent conditions at a threshold p-value of < 1 × 10^−04^. which is roughly comparable to the FDR 1 × 10^−04^. The retrieved gene set analysis was categorically arranged to eliminate redundancies. Categorization, previously described [25], was based on biological mapping pathways with a gamut from general pathways to specific pathways. For example, a general hallmark would be the immune pathway, where a more specific pathway would be cytokine signaling in immune systems, and a specific pathway would be interferon signaling. In regards to longevity gene data, a prior biological pathway list has been described [24]. Two independent individuals randomly collected, performed, and verified the analysis of categorization.

## 3. Results

### 3.1. Stress resistance

#### 3.1.1. Overview of stress resistance

Using more than 180 databases, we raked human genes relevant to stress resistance and identified 541 genes associated with the term, stress resistance, using the program, Genecards. The validation of the gene set was performed to run the gene set using another program, STRING-DB. Two pathway analyses showed categories relevant to stress resistance with the highest significance among all the pathways, the Reactome (FDR = 3.12 × 10^−51^) and the STRING-DB Biological process (FDR = 4.08 × 10^−59^). The results suggest that the gene set is enriched with the genes related to stress resistance. We found diverse ontology nomenclatures, including Biological, Molecular, Cellular, KEGG, and Reactome dimensions, among others, and thus we decided to use the Reactome analysis for consistency with our previous studies [24–25]. The gene set enrichment analysis for the 541 stress resistance genes compiled 398 biological Reactome pathways. Of them, 172 biological pathways met the threshold of a p-value < 1 × 10^−04^. The 172 biological pathways were summarized into 14 general pathways. General pathways, in the order of most frequent to least frequent, are: cell cycle, DNA repair, gene expression (transcription), metabolism of proteins, metabolism of RNA, signal transduction, cellular responses to stimuli, immune system, metabolism, autophagy, chromatin organization, developmental biology, organelle biogenesis, and DNA replication. Figure 1 summarizes the identified 14 general pathways and the sub-pathways with the highest number of hits for each general pathway.

**Figure 1.**
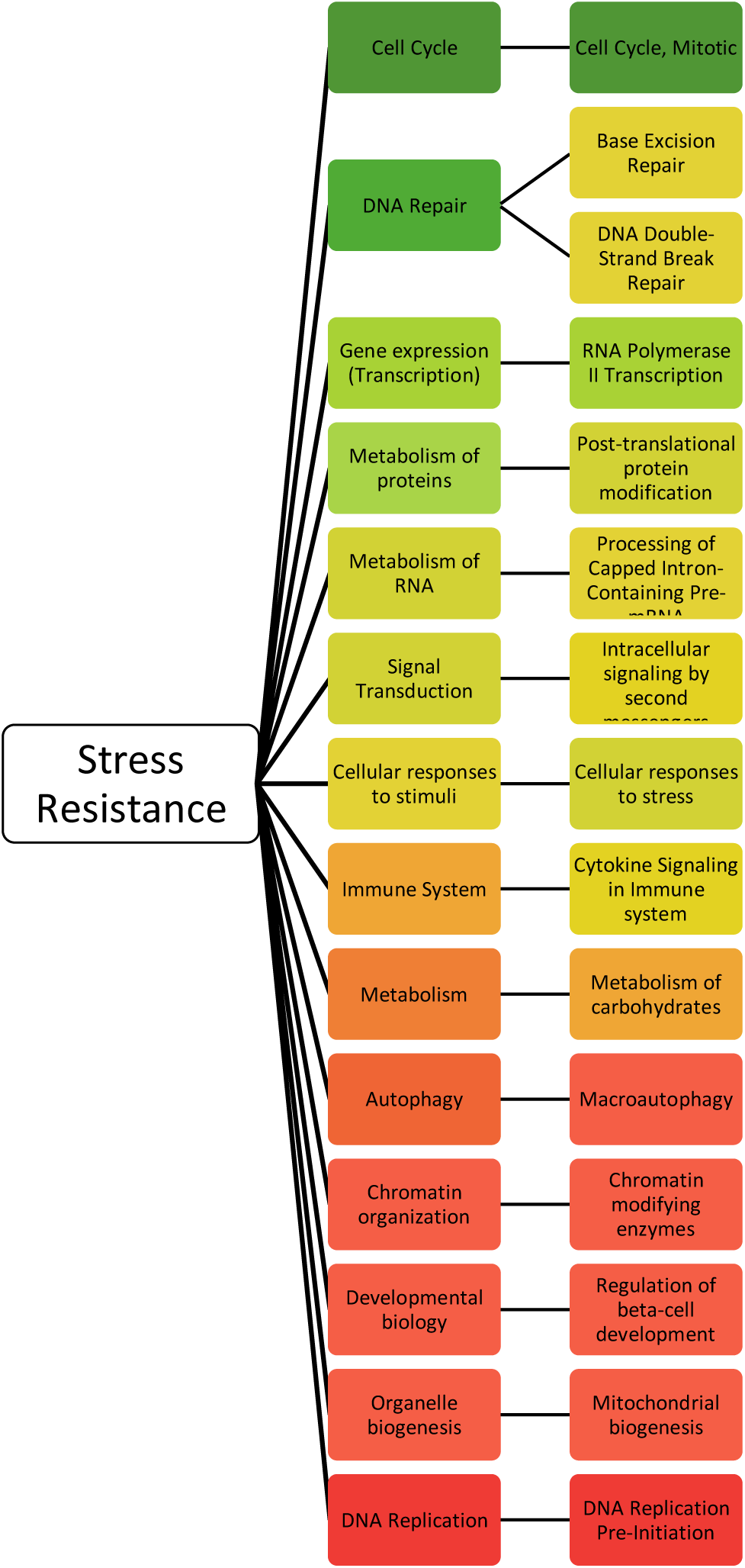
depicts the 14 general pathways linked to the stress resistance genes. The 14 general pathways are color-coded to the number of occurrences of the pathway. Green represents the general pathway with the most hits and red represents the general pathway with the least hits. To the right of the general pathways are the specific pathways. The specific pathways are color-coded to the number of occurrences of the pathway in relation to the other most frequent specific pathways.

#### 3.1.2. Stress resistance: Cell cycle-related pathways

Table 1 summarizes the results for the selected stress resistance biological pathways with the general pathway category cell cycle. Table 2 covers top hits and Table S1 covers all the 172 pathways. The cell cycle-related category was composed of four general pathways including cell cycle, DNA repair, DNA replication, and chromatin organization; they are related to the functions of the cell cycle (Table 1). The cell cycle preceded specific pathways in the order of the most frequent, M-phase, S-phase, and checkpoints (G1/S and mitotic spindle checkpoints). Notably, the M-phase pathways included specific phases of M-phase (prophase, metaphase, telophase, among others), sister chromatid cohesion and separation, meiosis, and nuclear envelope/lamina, including, for example, the Hutchinson-Gilford progeria syndrome (HGPS) gene (LMNA). Six of the cell cycle pathways included telomere maintenance and DNA synthesis, lagging and leading strands, and establishment of sister chromatid cohesion. The cellular response to stimuli included general response to stress, response to heat stress and HSF-1 regulations, and cellular senescence dependent on and independent of oxidative damage. The pathway category of chromatin organization included histone and modifiers, HDACs histone deacetylases, HDACs, and the NAD-dependent deacetylase sirtuin-1, SIRT1, which is involved in diverse functions of epigenetic regulations [27–28].

**Table 1.** Summary of stress resistance pathway categories identified by Reactome and the breakdowns.

| Pathways | No. of the pathways |
| --- | --- |
| **Total Number of Pathways Identified** | 172 |
| **Total Number of Pathway Categories Grouped\*** | 14 |

| Breakdowns of 172 Pathway categories | No. of the pathways |
| --- | --- |
| 1 Cell Cycle | 40 |
| 2 DNA Repair | 37 |
| 3 Gene expression (Transcription) | 19 |
| 4 Metabolism of proteins | 17 |
| 5 Metabolism of RNA | 14 |
| 6 Signal Transduction | 14 |
| 7 Cellular responses to stimuli | 11 |
| 8 Immune System | 6 |
| 9 Metabolism | 4 |
| 10 Autophagy | 3 |
| 11 Chromatin organization | 2 |
| 12 Developmental biology | 2 |
| 13 Organelle biogenesis | 2 |
| 14 DNA Replication | 1 |
| Total | 172 |

**Table 2.**
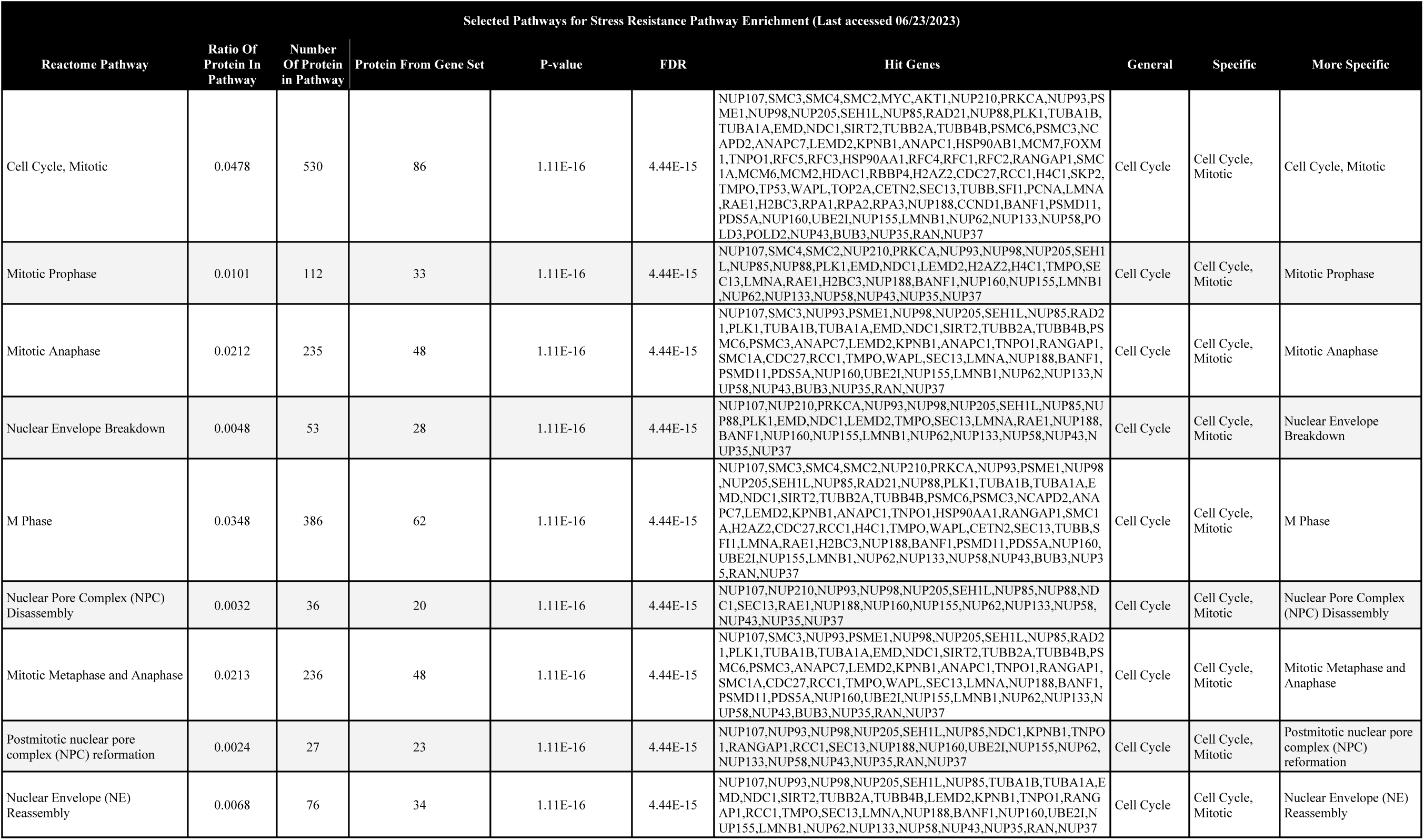
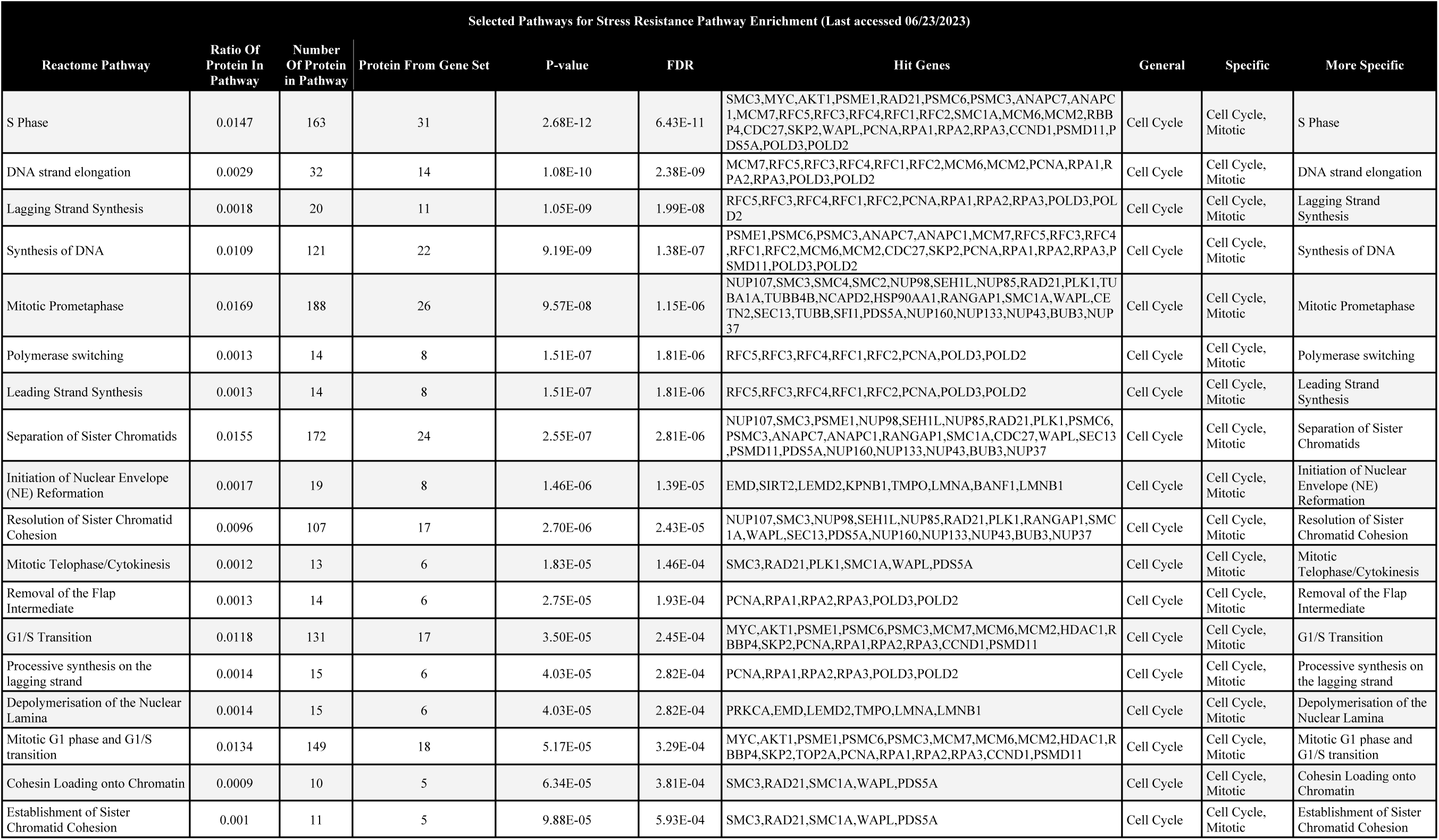
Top 27 stress-resistance pathways. The general, specific, and more specific pathways were organized and listed for each of the 27 Reactome pathways. All of the pathways in Table 1 fall under the cell cycle being the overarching pathway. The ratio of protein in the pathway, number of proteins in the pathway, protein from the gene set, p-value, FDR, and hit genes were provided by executing a gene set enrichment analysis.

The second-most frequent hit of the 14 general pathways was within the category of DNA repair. DNA repair is known to link to cell cycle functions including DNA replication and checkpoint. The most frequent specific pathways for DNA Repair covered most if not all repair systems, including base excision repair and DNA double-strand break repair each with 9 hits, while others included DNA damage bypass (8 pathways), mismatch repair (3 pathways), and nucleotide excision repair (8 pathways). Importantly, the pathways included another progeria gene, the Werner syndrome gene (WRN), which is a RecQ-like DNA helicase gene. WRN was involved in the cell cycle, such as cell cycle checkpoints, and telomere maintenance; in the DNA-related pathways, such as homology-directed repair, and double-strand break repair; and the gene expression.

#### 3.1.3. Stress resistance: Signal transduction-gene expression pathways

The signal transduction-related pathways included two major pathway categories, signal transduction and gene expression. The signal transduction pathways identified 14 pathways, composed of intracellular signaling by second messengers and other signaling pathways. They included stress-resistance pathways, which can be divided into several known pathways mediated by insulin-PIP3-AKT-FOXO, NOTCH, nuclear receptors (ESR-mediated estrogen-dependent gene expression), Rho GTPases, and WNT-beta-catenin). Importantly, mTOR was involved in and scattered among 13 pathways ranging from macroautophagy (1 pathway), cellular stress response (4 pathways), RNA Polymerase II Transcription (2 pathways), regulation of p53 (2 pathways), and PIP3/PTEN-AKT pathways (4 pathways).

The gene-expression pathways were identified as 19 pathways, including RNA polymerase II transcription (15 specific pathways), epigenetic regulation of gene expression (2 specific pathways), and gene silencing by RNA (2 specific pathways). RNA polymerase II transcription included the notable genes, TP53, FOXO (FOXO1,3,4), RUNX2, VENTX, Notch-HLH, p14^ARF^ among others. Epigenetic regulation of gene expression included ERCC6, EHMT2, HDACs (HDAC1 and 2), and histone genes (H2s and H4C1). Gene silencing by RNA included a total of 25 genes, including mostly NUPs and histone genes (H2AZ2 and H4C1). Interestingly, SIRTs (SIRT1,3), known as gene silencing genes, were identified as genes in gene expression but not as those in epigenetic regulation of gene expression.

#### 3.1.4. Stress resistance: Metabolic pathways

The metabolic pathways included the metabolism of carbohydrates, the metabolism of non-coding RNA, and organelle biogenesis and maintenance, according to the Reactome categories. The categories, metabolism of proteins and metabolism of RNA, are described in Section 3.1.5. Of 172 pathways, 4 pathways fell into the metabolism of carbohydrates, while 2 pathways belong to organelle biogenesis and maintenance in the mitochondria (Table S1). Interestingly, the metabolism of carbohydrates included mostly proteins in the transport and regulation of metabolic enzymes, including the nuclear pore proteins, nucleoporin (NUP) genes (16 genes), and glucokinase which involves transport systems between the cytoplasm and the nucleus. Although the vast majority of the genes in this group fell into the regulatory genes of metabolism, they were mostly not in the metabolic enzymes. Thus, this category represented the transport and regulation of them rather than the metabolic enzymes themselves. Importantly, the nuclear pore proteins (NUPs) were enriched in 46 biological pathways relevant to the transport of macromolecules and nuclear envelope assembly; they were involved in glucose and carbohydrate metabolisms (4 pathways), pre-matured and matured RNA (11 pathways), SUMOylation-dependent protein degradation (9 pathways), cellular response to heat (2 pathways), interferon anti-virus immune system (3 pathways), cell cycle (14 pathways), and gene silencing (2 pathways), among others. The result highlights the role of the nuclear envelope integrity and the transport of macromolecules in the biological pathways for stress resistance. The result also suggests that the category of metabolism may not represent metabolic enzymes but mostly proteins in the transport and regulation of metabolism. Importantly, the accelerated aging syndrome (progeria) gene, HGPS gene (LMNA) is also included in this group related to the structure of the nuclear envelope and pores.

#### 3.1.5. Stress resistance: Metabolism of proteins, metabolism of RNA and others

Metabolism of proteins: Of 172 pathways, 11 pathways were identified as the category metabolism of proteins, all of which were proteins involved in protein degradation called SUMOylation. They include SUMOylation of ubiquitinated proteins, chromatin organization proteins, DNA damage response and repair proteins, SUMOylation proteins, RNA binding proteins, and DNA replication proteins. Thus, they overlapped with the cell cycle (DNA repair, replication, and chromatin organizations). As described above, 9 out of 11 pathways were enriched with the nuclear pore proteins (NUPs).

Other categories were as follows. Mitochondrial biogenesis: 2 pathways were identified as the mitochondrial biogenesis pathways, which were a part of energy metabolism. Metabolism of RNA: 9 pathways were identified as the Metabolism of RNA pathways. They were involved in the processing of capped intron-containing pre-mRNA including the transport of mature mRNAs derived from transcripts (7 pathways) and mRNA splicing (2 pathways). Cellular responses to stimuli: 11 pathways fell into this category, including heat response dependent on HSF-1 (5 pathways) and HSP90 chaperon (1 pathway), response to starvation (1pathway), response to starvation (1 pathway), response to amino acid deficiency (1 pathway), cellular senescence (2 pathways) including senescence induced by oxidative stress, among others. Immune system: 6 pathways, including innate immunity (1 pathway), and cytokine signaling (5 pathways) such as interferon signaling (3 pathways) and interleukin signaling (1 pathway). Autophagy: 3 pathways included macroautophagy (2 pathways) and chaperone-mediated autophagy (1 pathway). Developmental biology: 2 pathways identified beta-cell development, which is essential for insulin secretion.

### 3.2. Longevity

The 357 longevity genes and the detailed pathways have been described previously [24]. Top-hit longevity pathways are shown in Table 3. In this study, we performed a more in-depth analysis of the pathways with a threshold of a p-value < 1 × 10^−04^. Firstly, the gene set enrichment analysis identified 141 biological pathways, narrowed down to 50 biological pathways with the p-value threshold. The 50 biological pathways were summarized into 7 general pathways. The 7 general pathways, in the order of most frequent to least frequent result, are gene expression (transcription), signal transduction, immune system, cellular responses to stimuli, metabolism, transport of small molecules, and autophagy. Figure 2 summarizes the identified 7 general pathways and the most frequent sub-pathways for each of the 7 general pathways.

**Figure 2.**
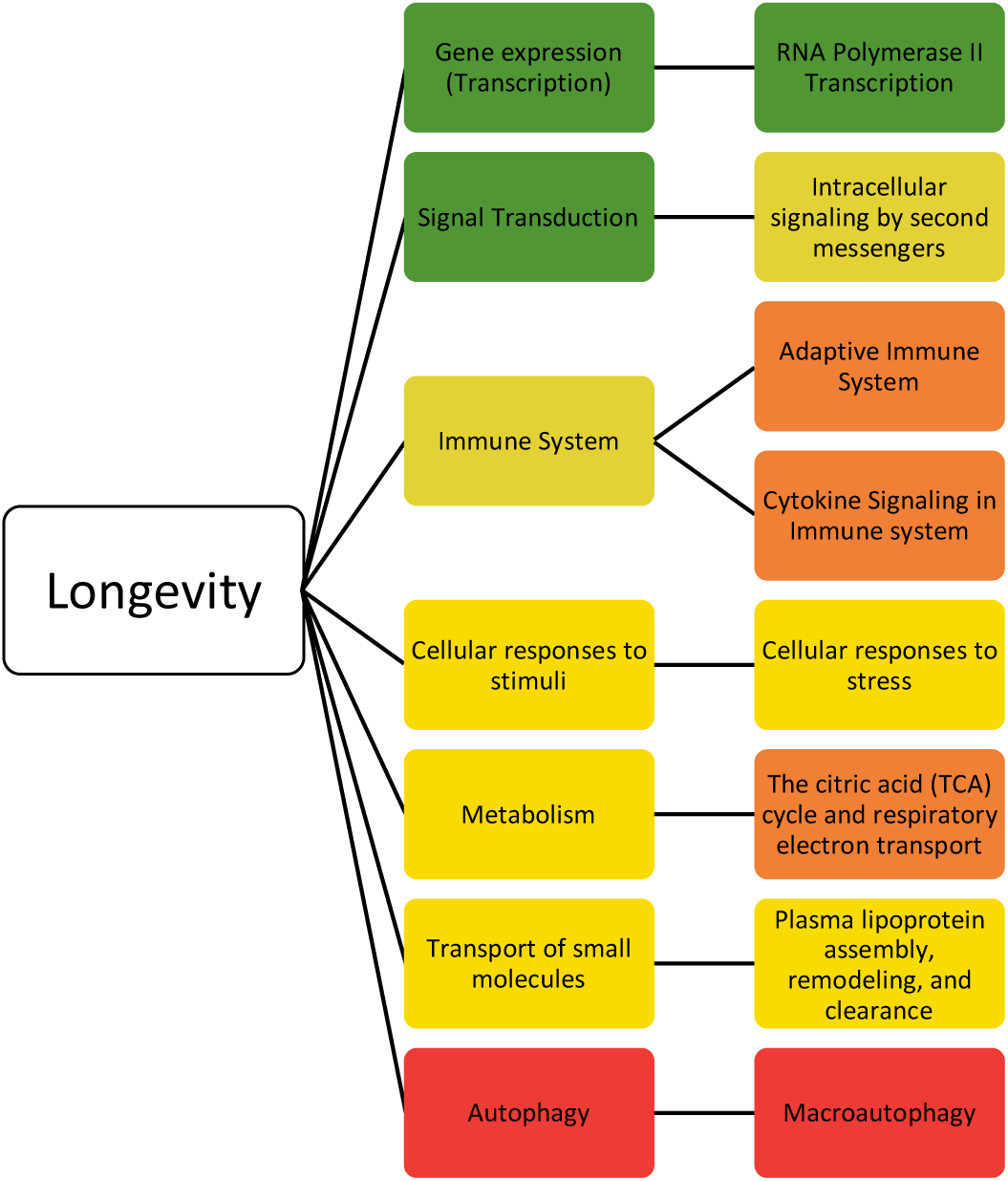
depicts the 7 general pathways associated with the longevity genes. The color of the general pathway represents the number of occurrences of the pathway relative to the other general pathways. The pathways in green have the greatest number of hits whereas the red have the fewest number of hits. To the right of the general pathways are the most frequent specific pathways. The specific pathways are color-coded in relation to the other most frequent specific pathways with green being the most frequent and red being the least frequent.

**Table 3.**
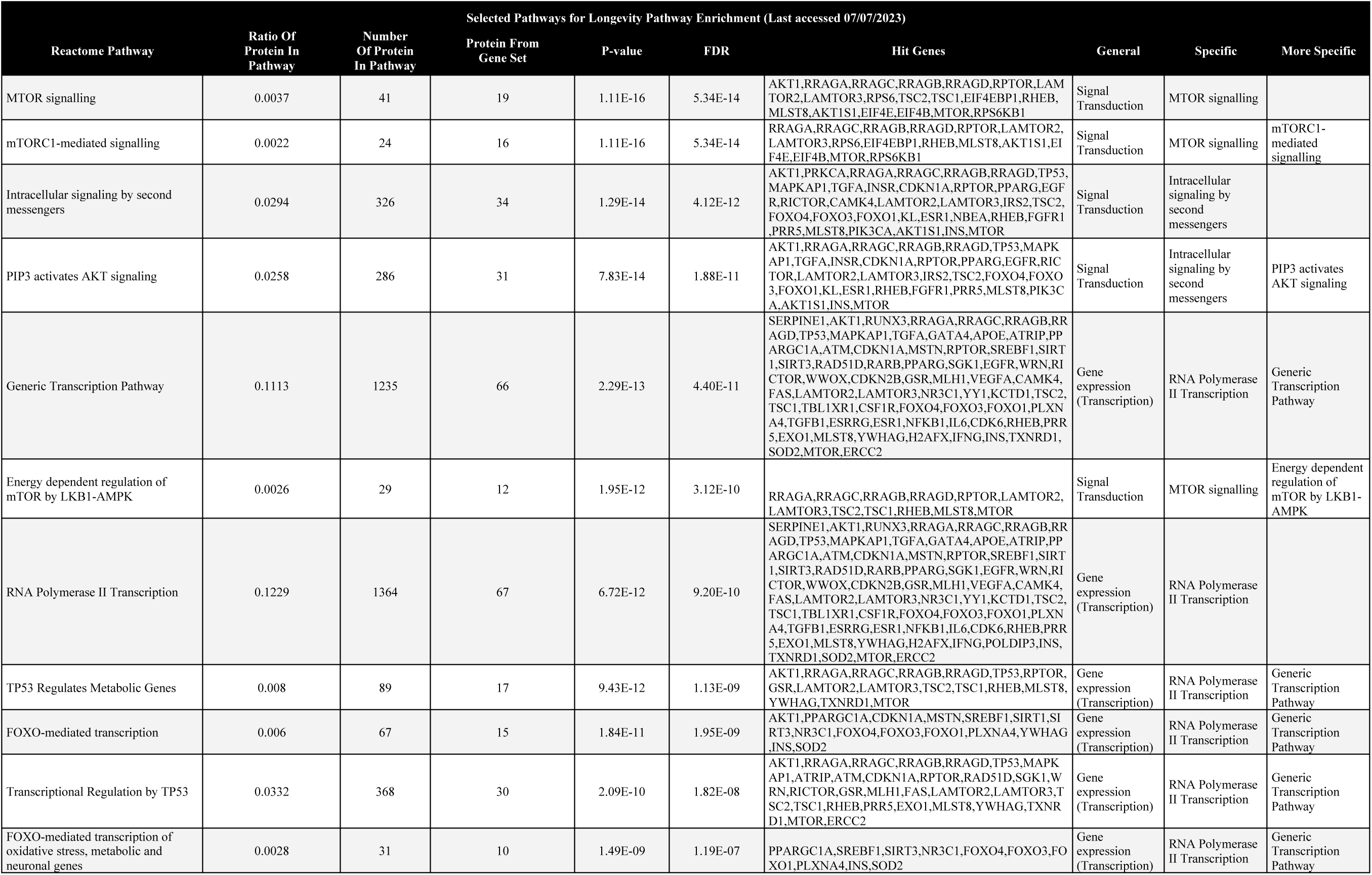
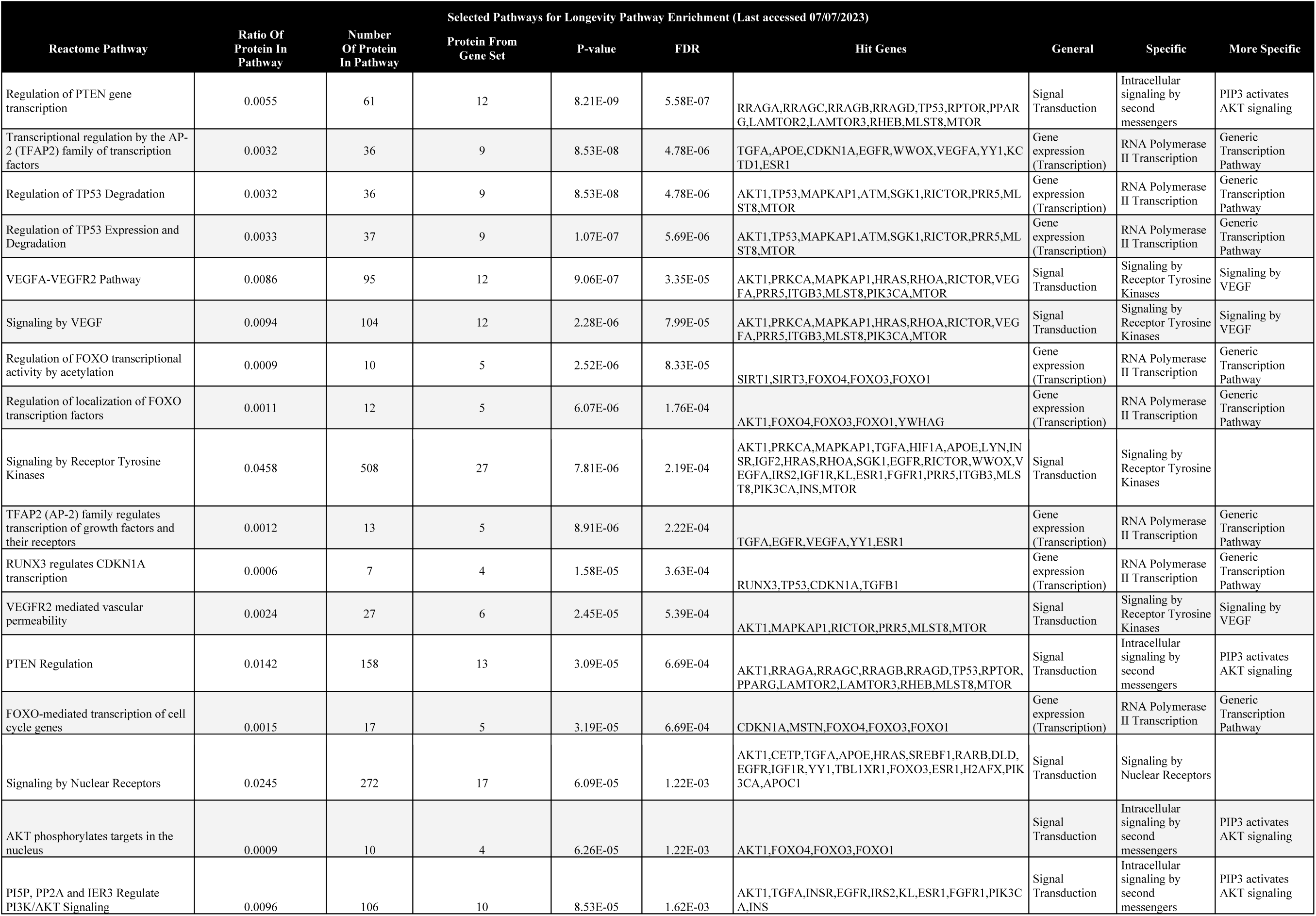
28 top-hit longevity pathways were shown. The general, specific, and more pathways were traced for the Reactome pathways. All of the pathways in Table 3 fall under the gene expression (transcription) or signal transduction pathways.

Out of 50 pathways, the highest number of hits fell into gene expression (transcription) and signal transduction with 14 pathways each. As expected, the pathways including FOXO, which are known to affect lifespans in the model systems, are overlapped between gene expression and signal transduction: 7 out of 14 pathways as gene expression (FOXO-mediated gene expression) and 5 pathways as signal transduction (PIP3/PTEN-AKT pathways). The remaining 7 gene-expression pathways fell into TP53 regulation (4 pathways), AP-2 (TFAR2) regulation (2 pathways), and RUNX3-dependent CDKN1A transcription (1 pathway). Another hit among the more specific pathways within signal transduction was intracellular signaling by second messengers. Table 2 summarizes the results of the selected longevity biological pathways with gene expression (transcription) and signal transduction as the general pathways.

### 3.3 Overlap for Stress Resistance and Longevity

Figure 3 summarizes biological pathway categories overlapped and non-overlapped between stress resistance and longevity. 6 pathway categories overlapped between the stress resistance genes and longevity genes. The overlapping general pathways were gene expression (transcription), signal transduction, cellular responses to stimuli, immune system, metabolism, and autophagy. Importantly, the overlapped categories include the PIP3-AKT-FOXO and mTOR pathways (categories: gene expression and signal transduction) (Table S2), which are known to control life extension and/or stress resistance in the model systems. They also included the accelerated aging syndrome (progeria) genes, including the Hutchinson-Gilford progeria syndrome (HGPS) gene (LMNA) and Werner syndrome gene (WRN) (categories: cellular response to stimuli/stress), while the genes were also included in non-overlapped categories (DNA repair and cell cycle) (Table S2). The other longevity pathways included the transport of small molecules, which were involved in lipoprotein metabolism. Notably, dysregulation of lipoprotein metabolism (i.e., oxidation and glycation) can cause atherosclerotic lesions, which, in broader classification, may be considered as stress damage. In summary, the longevity pathways mostly overlapped with stress resistance pathways. In contrast, the other stress-resistance pathways not overlapped in longevity were specific to molecular (DNA and RNA) and biological categories (cell cycle, chromatin, organelle, post-translational modification/metabolism of proteins) (Figure 3).

**Figure 3.**
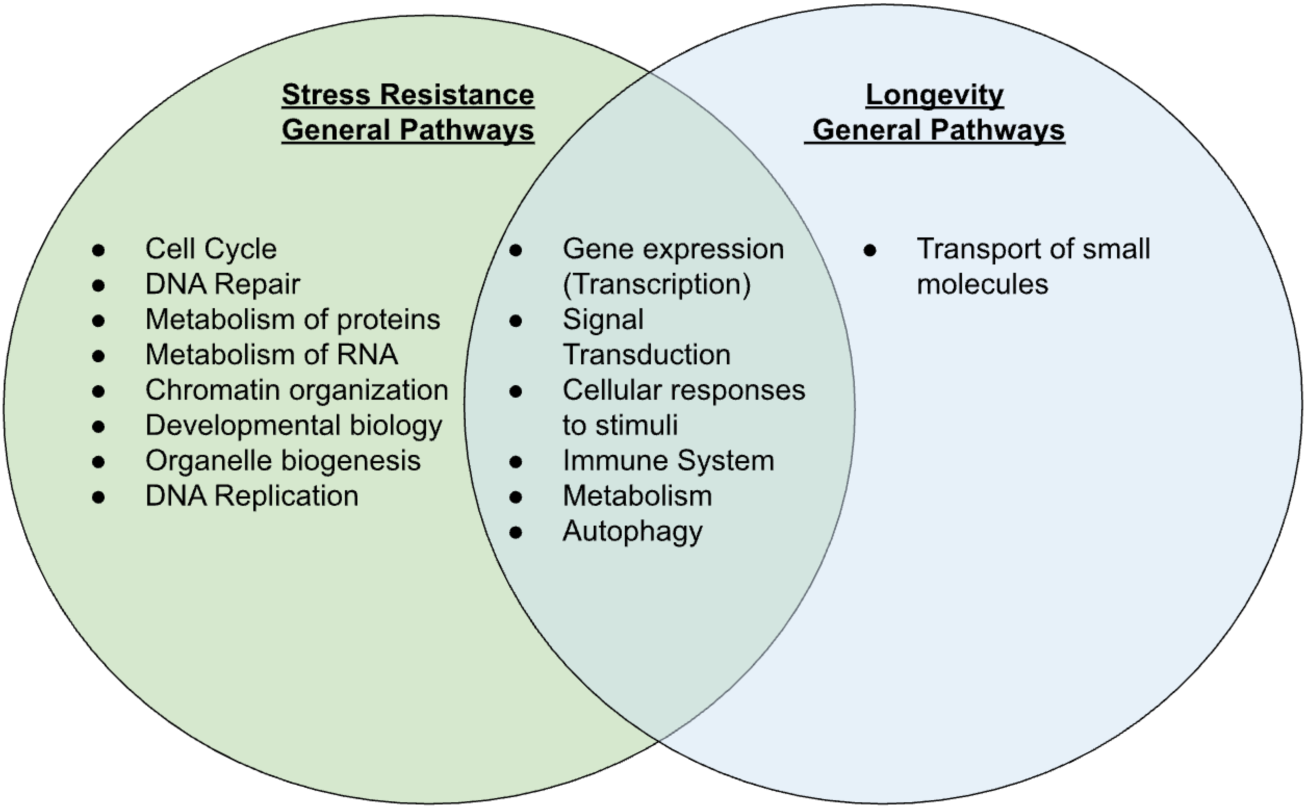
compares and contrasts the general pathways linked to the stress resistance genes and longevity genes. 8 general pathways are linked to only stress resistance whereas 1 general pathway is linked to only longevity. 6 general pathways are linked to both stress resistance and longevity.

## 4. Discussion

In this study, we used gene set enrichment analysis (GSEA) to uncover the gene networks to explore the gene networks associated with stress resistance and longevity in humans. We successfully identified 541 genes linked to stress resistance, which is more comprehensive than the previously documented 215 single nucleotide polymorphisms associated with longevity in individuals who smoke [29]. The results show that the genes are involved in 172 pathways. Moreover, we found that these pathways can be grouped into 14 major pathways, closely related to six out of the seven longevity pathways previously obtained from 50 biological pathways.

### 4.1. Similarity among stress resistance and longevity pathways

Six pathway categories are common between stress resistance and longevity genes. The categories include gene expression (transcription), signal transduction, immune system, cellular responses to stimuli, metabolism, and autophagy. In-depth investigation identifies the well-known pathways for stress resistance and longevity in the model systems, such as the PIP3/PTEN-AKT-FOXO pathway that are covered mainly under the gene expression and the signal transduction pathways. These pathways are regulatory that govern other biological pathways. If life extension is the result of reducing the causes of aging, then stress resistance should be involved in the causes of aging (i.e., stress resistance related to longevity). Importantly, the common categories also include the progeria genes (HGPS/LMNA and WRN), while the genes are also included in non-overlapped categories.

The common pathway categories between stress resistance and longevity suggest underlying mechanisms. Stress resistance confers increased survival after exposure to oxidative stress, heat, UV light, DNA-damaging agents, toxic peptides, and other types of stress. Previous studies suggest that multiplex stress resistance unifies multiple forms of single stress resistance and is essential for life extension [13,16]. It seems reasonable to state that a wide variety of stress resistance is under the control of limited biological mechanisms, for example, serotonin, insulin/IGF-1 pathways [20–23,30] as well as the genetic mechanisms for stress-resistance categories related to longevity. It is worth noting that stress resistance may be altered depending on the intrinsic stress conditions, for example, supplementing glucose metabolites (pyruvate and lactate) causes increases in oxidative stress, and triggers hormesis (also called biological resilience) and life extension [31].

### 4.2. Differences among stress resistance and longevity pathways

In this study, a clear difference between stress resistance and longevity is that stress resistance covers broader pathways; 172 stress-resistance pathways are summarized into 14 categories, while 141 pathways are summarized into 7 categories. Of 14, eight stress resistance categories unique to stress are enriched in the functions related to cell proliferation and development, including cell cycle, DNA repair, metabolism of proteins, metabolism of RNA, chromatin organization, developmental biology (of beta-cell), organelle biogenesis (in the mitochondria), and DNA replication.

There are two major differences between experimental stress resistance and longevity: (1) nature of stress damage: Stress resistance assesses survival under the exposure to a sublethal stressor, while such excess stress is not expected in normal aging; (2) sites of stress damage: each form of excess stress has a specific site of damage in the whole body, while any stress is experimentally minimized in normal aging. Thus, single stress resistance is segmental and may be insufficient for life extension. Resistance to single stress (e.g., oxidative stress and heat) can be separated from life extension in worms [8,32]. Importantly, suppressing superoxide dismutase genes reduces life extension by a life-extending *daf-2* (Insulin/IGF-1 receptor gene) mutation [15, 32], suggesting that single stress resistance may still count for a part of life extension.

Notably, lipoprotein metabolism is included in longevity but not stress resistance in the Reactome-defined pathways. However, low-density lipoprotein (LDL) is involved in a part of lipoprotein metabolism, which is known to trigger atherosclerotic lesions by oxidation and glycation [33–34], and thus, may be viewed as stress damage on LDL. Thus, we propose to include a Reactome subcategory of stress damage under the category of lipoprotein metabolism. In humans, the leading cause of death has been cardiovascular diseases [35–36]. This suggests specific parts of the body, such as the circulation systems, play a critical role in mortality.

### 4.3. Advantages and limitations of the method

The advantages and limitations of the Reactome analysis have been discussed previously [24]. Human phenotypes primarily result from multiple polymorphisms and multifactorial differences, including genetic, phenotypic, and clinical heterogeneity among others [24–25,37–40]. The gene-network analysis, such as GSEA, effectively explores conditions in humans in which single-gene mutational approaches are not impractical. Visualizing the genetic network based on the pathway categories provides insights that may have been missed previously. Considering that most gene effects in humans are epistatic, quantitative, or multifactorial, gene network analysis has a valuable means of overviewing the genetic effects in humans.

The limitations of GSEA are as follows. Firstly, the list of the genes generated is a living knowledge that needs to be updated periodically. Secondly, the analysis outputs are multi-dimensional ontology pathways organized by each dimension, ranging from molecules, cellular/extracellular, tissue, functional, and phenotypic among others, where nomenclature is different to indicate similar biological phenomena. For example, we found the pathway categories, cellular response to cells, in the Reactome analysis and STRING-DB biological pathways; however, other pathway analyses are not straightforward to identify the categories related to stress resistance.

Moreover, we had difficulty with terminology aligning with the aging hallmarks previously reported [41], which are defined as a mixture of various ontology dimensions ranging from biological and other categories, for example, categories related to nutrition (deregulated nutrient-sensing), stem cell (stem cell exhaustion), and telomere (telomere attrition), are at the levels of the macromolecule, the cell, and the chromosomal domain, which are at the different ontology dimensions. It is also not clear how the aging hallmarks fit in well-established pathways for longevity and stress resistance, including the Insulin/IGF-1 pathways, whose second messengers are PIP3-AKT-FOXO, and the mTOR pathways. To minimize discrepancies and have more consistency, we have used the Reactome analysis, which has been relatively accepted and incorporated into various network analyses. Finally, due to the reason above, the aging hallmark terminologies do not align with the ontology pathway categories with various dimensions. We propose to have more precise biological terminologies defining multi-dimensional nomenclature, including molecular, cellular/extracellular, tissue, and systemic levels among others so that it would be more unified and integral knowledge as the new findings keep coming into this field.

### 4.4 Nuclear transport and integrity emphasized stress resistance and aging but not longevity

We found that nuclear pore proteins (NUPs) are enriched in 46 biological pathways for stress resistance (Table S1). NUPs are relevant to the transportation of macromolecules and the assembly of the nuclear envelope. NUPs are the main components of the nuclear pore complexes. They are associated with scaffold complexes including the NUP107-160 complex and NUP93-205 complex, the NUP214 complex, the NUP98 complex, and the NUP62 complex, among others [42–43] (D’Angelo and Hetzer, 2008; Martins et al., 2020). The NUP62, 93, and 152 play a role in nuclear transport, which is suggested to be involved in aging [43]. Additionally, nuclear envelope integrity has been proposed to play a role in aging. Firstly, the accelerated aging syndrome (progeria), the Hutchinson-Gilford progeria syndrome (HGPS) gene is, caused by mutations in the LMNA gene that encodes Lamin A/C [44] and it is tightly related to the structure of the nuclear envelope and pores [43,45]. In this study, the NUPs are involved in diverse pathways including carbohydrate metabolism of carbohydrates, RNA and proteins, cellular stress response, anti-virus immune system, cell cycle, and gene silencing, among others. However, the NUPs are not involved in longevity pathways. We suggest that the NUP-dependent nuclear integrity and transport play a role in aging and stress resistance but not longevity. Taken together, the results suggest a novel role of the NUP-dependent nuclear integrity and transport in age-related metabolic changes and diverse pathways.

## 5. Conclusions

This study indicates that the biological pathways responsible for longevity are closely linked with those involved in stress resistance. The finding suggests that the pathways that play a role in longevity are also important for stress resistance, which aligns with previous studies on multiplex stress resistance [5,16–18]. We found that longevity pathways were summarized into six categories. Assuming that suppressing the causes of aging should delay aging and lead to life extension, the causes of aging may be limited. Stress resistance related to longevity should be involved in the causes of aging, while stress resistance unrelated to longevity may be the consequence of aging, responding to stress damage. Similarly, progeria genes are involved in DNA and nuclear integrity [46–47], which may be involved in both the cause and consequences of aging. The findings in this study appear to support initial age-related causes transitioning to more complex consequences of aging with a mixture of aging physiology and pathology. Previous studies suggest that there is a transitional phase before advanced aging; the transitional phase formulated the ground of the aging theory (middle-life crisis theory) [22–23].

This study updates that multiplex stress resistance involves mechanisms unifying responses and resistances to multiple forms of stress. In addition, stress resistance is likely not limited to molecular damage but also extends to lipid/lipoprotein metabolism that includes oxidation and glycation of LDL not currently classified as damage in the Reactome ontology analysis. This study contributes to a critical step forward in the fight against aging and age-related diseases. This study reinstates the significance of stress resistance and longevity, while it supports distinct mechanisms of aging and longevity.

## Author Contributions

Data generation, analysis, validation, and initial manuscript preparation and editing, K.B., P.L., A.S.; all aspects of the research project, S.M. All authors have read and agreed to the published version of the manuscript.

## Institutional Review Board Statement

## Informed Consent Statement

## Data Availability Statement

All the data has been included in the manuscript.

## Acknowledgments

We gratefully appreciate Amy Sakazaki, Haider Shah, and other members of Murakami Laboratory for the initial technical assistance and useful discussion. The manuscript was proofread and revisions were made for clarity and readability by the members and by grammar check software embedded with AI (https://app.grammarly.com).

## References

1. Harman D. Aging: a theory based on free radical and radiation chemistry. J Gerontol. 1956 11(3):298–300. doi: 10.1093/geronj/11.3.298.

2. Muller FL, Lustgarten MS, Jang Y, Richardson A, Van Remmen H. Trends in oxidative aging theories. Free Radic Biol Med. 2007 43(4):477–503. doi: 10.1016/j.freeradbiomed.2007.03.034.

3. Doonan R, McElwee JJ, Matthijssens F, Walker GA, Houthoofd K, Back P, Matscheski A, Vanfleteren JR, Gems D. Against the oxidative damage theory of aging: superoxide dismutases protect against oxidative stress but have little or no effect on life span in Caenorhabditis elegans. Genes Dev. 2008 22(23):3236–41. doi: 10.1101/gad.504808.

4. Pérez VI, Bokov A, Van Remmen H, Mele J, Ran Q, Ikeno Y, Richardson A. Is the oxidative stress theory of aging dead? Biochim Biophys Acta. 2009 1790(10):1005–14. doi: 10.1016/j.bbagen.2009.06.003.

5. Dues DJ, Andrews EK, Schaar CE, Bergsma AL, Senchuk MM, Van Raamsdonk JM. Aging causes decreased resistance to multiple stresses and a failure to activate specific stress response pathways. Aging (Albany NY). 2016 8(4):777–95. doi: 10.18632/aging.100939.

6. Salmon AB, Richardson A, Pérez VI. Update on the oxidative stress theory of aging: does oxidative stress play a role in aging or healthy aging? Free Radic Biol Med. 2010 Mar 1;48(5):642–55. doi: 10.1016/j.freeradbiomed.2009.12.015.

7. Ziada AS, Smith MR, Côté HCF. Updating the Free Radical Theory of Aging. Front Cell Dev Biol. 2020 8:575645. doi: 10.3389/fcell.2020.575645.

8. Douglas PM, Baird NA, Simic MS, Uhlein S, McCormick MA, Wolff SC, Kennedy BK, Dillin A. Heterotypic Signals from Neural HSF-1 Separate Thermotolerance from Longevity. Cell Rep. 2015 Aug 18;12(7):1196–1204. doi: 10.1016/j.celrep.2015.07.026. Epub 2015 Aug 6. PMID: 26257177; PMCID: PMC4889220.

9. Wang J, Robida-Stubbs S, Tullet JM, Rual JF, Vidal M, Blackwell TK. RNAi screening implicates a SKN-1-dependent transcriptional response in stress resistance and longevity deriving from translation inhibition. PLoS Genet. 2010 6(8):e1001048. doi: 10.1371/journal.pgen.1001048.

10. Zhang P, Zhai Y, Cregg J, Ang KK, Arkin M, Kenyon C. Stress Resistance Screen in a Human Primary Cell Line Identifies Small Molecules That Affect Aging Pathways and Extend *Caenorhabditis elegans*’ Lifespan. G3 (Bethesda). 2020 10(2):849–862. doi: 10.1534/g3.119.400618.

11. Derisbourg MJ, Wester LE, Baddi R, Denzel MS. Mutagenesis screen uncovers lifespan extension through integrated stress response inhibition without reduced mRNA translation. Nat Commun. 2021 12(1):1678. doi: 10.1038/s41467-021-21743-x.

12. Harper JM, Salmon AB, Chang Y, Bonkowski M, Bartke A, Miller RA. Stress resistance and aging: influence of genes and nutrition. Mech Ageing Dev. 2006 127(8):687–94. doi: 10.1016/j.mad.2006.04.002.

13. Murakami S. Stress resistance in long-lived mouse models. Exp Gerontol. 2006 Oct;41(10):1014–9. doi: 10.1016/j.exger.2006.06.061.

14. Zhou KI, Pincus Z, Slack FJ. Longevity and stress in Caenorhabditis elegans. Aging (Albany NY). 2011 3(8):733–53. doi: 10.18632/aging.100367.

15. Soo SK, Rudich ZD, Ko B, Moldakozhayev A, AlOkda A, Van Raamsdonk JM. Biological resilience and aging: Activation of stress response pathways contributes to lifespan extension. Ageing Res Rev. 2023 88:101941. doi: 10.1016/j.arr.2023.101941.

16. Murakami S, Johnson TE. A genetic pathway conferring life extension and resistance to UV stress in Caenorhabditis elegans. Genetics. 1996 143(3):1207–18. doi: 10.1093/genetics/143.3.1207.

17. Murakami S, Salmon A, Miller RA. Multiplex stress resistance in cells from long-lived dwarf mice. FASEB J. 2003 17(11):1565–6. doi: 10.1096/fj.02-1092fje.

18. Salmon AB, Murakami S, Bartke A, Kopchick J, Yasumura K, Miller RA. Fibroblast cell lines from young adult mice of long-lived mutant strains are resistant to multiple forms of stress. Am J Physiol Endocrinol Metab. 2005 289(1):E23–9. doi: 10.1152/ajpendo.00575.2004.

19. Dues DJ, Andrews EK, Schaar CE, Bergsma AL, Senchuk MM, Van Raamsdonk JM. Aging causes decreased resistance to multiple stresses and a failure to activate specific stress response pathways. Aging (Albany NY). 2016 8:777–795. doi: 10.18632/aging.100939

20. Murakami H, Murakami S. Serotonin receptors antagonistically modulate Caenorhabditis elegans longevity. Aging Cell. 2007 6(4):483–8. doi: 10.1111/j.1474-9726.2007.00303.x.

21. Murakami H, Bessinger K, Hellmann J, Murakami S. Manipulation of serotonin signal suppresses early phase of behavioral aging in Caenorhabditis elegans. Neurobiol Aging. 2008 29(7):1093–100. doi: 10.1016/j.neurobiolaging.2007.01.013.

22. Murakami, S., Cabana, K. and Anderson, D. Current Advances in the Studies of Oxidative Stress and Age-Related Memory Impairment in *C. elegans*. In Oxidative Stress in Vertebrates and Invertebrates (eds T. Farooqui and A.A. Farooqui). 2011 10.1002/9781118148143.ch25

23. Murakami, S. Age-dependent modulation of learning and memory in *Caenorhabditis elegans*, in Invertebrate learning and memory; handbook of behavioral neuroscience, eds R. Menzel and P. R. Benjamin (Cambridge, MA: Elsevier/Academic Press), 2013 pp 140–150. doi: 10.1016/B978-0-12-415823-8.00012-5

24. Balmorez T, Sakazaki A, Murakami S. Genetic Networks of Alzheimer’s Disease, Aging, and Longevity in Humans. Int J Mol Sci. 2023 24(6):5178. doi: 10.3390/ijms24065178.

25. Murakami S, Lacayo P. Biological and disease hallmarks of Alzheimer’s disease defined by Alzheimer’s disease genes. Front Aging Neurosci. 2022 14:996030. doi: 10.3389/fnagi.2022.996030.

26. Stelzer G, Rosen N, Plaschkes I, Zimmerman S, Twik M, Fishilevich S, Stein TI, Nudel R, Lieder I, Mazor Y, Kaplan S, Dahary D, Warshawsky D, Guan-Golan Y, Kohn A, Rappaport N, Safran M, Lancet D. The GeneCards Suite: From Gene Data Mining to Disease Genome Sequence Analyses. Curr Protoc Bioinformatics. 2016 54:1.30.1–1.30.33. doi: 10.1002/cpbi.5.

27. Stünkel W, Campbell RM. Sirtuin 1 (SIRT1): the misunderstood HDAC. J Biomol Screen. 2011 16(10):1153–69. doi: 10.1177/1087057111422103.

28. Alves-Fernandes DK, Jasiulionis MG. The Role of SIRT1 on DNA Damage Response and Epigenetic Alterations in Cancer. Int J Mol Sci. 2019 Jun 28;20(13):3153. doi: 10.3390/ijms20133153.

29. Levine ME, Crimmins EM. A Genetic Network Associated With Stress Resistance, Longevity, and Cancer in Humans. J Gerontol A Biol Sci Med Sci. 2016 71(6):703–12. doi: 10.1093/gerona/glv141.

30. Machino K, Link CD, Wang S, Murakami H, Murakami S. A semi-automated motion-tracking analysis of locomotion speed in the C. elegans transgenics overexpressing beta-amyloid in neurons. Front Genet. 2014 5:202. doi: 10.3389/fgene.2014.00202.

31. Tauffenberger A, Fiumelli H, Almustafa S, Magistretti PJ. Lactate and pyruvate promote oxidative stress resistance through hormetic ROS signaling. Cell Death Dis. 2019 10(9):653. doi: 10.1038/s41419-019-1877-6.

32. Dues DJ, Andrews EK, Senchuk MM, Van Raamsdonk JM. Resistance to Stress Can Be Experimentally Dissociated From Longevity. J Gerontol A Biol Sci Med Sci. 2019 12;74(8):1206–1214. doi: 10.1093/gerona/gly213.

33. Petrucci G, Rizzi A, Hatem D, Tosti G, Rocca B, Pitocco D. Role of Oxidative Stress in the Pathogenesis of Atherothrombotic Diseases. Antioxidants (Basel). 2022 Jul 20;11(7):1408. doi: 10.3390/antiox11071408. PMID: 35883899; PMCID: PMC9312358.

34. Lyons TJ. Glycation and oxidation: a role in the pathogenesis of atherosclerosis. Am J Cardiol. 1993 Feb 25;71(6):26B–31B. doi: 10.1016/0002-9149(93)90142-y. PMID: 8434558.

35. WHO, Global Health Estimates: Life expectancy and leading causes of death and disability (2020) https://www.who.int/data/gho/data/themes/mortality-and-global-health-estimates (Last accessed on July 13, 2023)

36. CDC, Mortality in the United States, 2022, In Deaths and Mortality, National Center for Health Statistics, Available Online: https://www.cdc.gov/nchs/fastats/deaths.htm (last accessed on July 13, 2023)

37. Mackay TF. Epistasis and quantitative traits: using model organisms to study gene-gene interactions. Nat Rev Genet. 2014 15(1):22–33. doi: 10.1038/nrg3627.

38. Mackay TF, Moore JH. Why epistasis is important for tackling complex human disease genetics. Genome Med. 2014 6(6):124. doi: 10.1186/gm561. Erratum in: Genome Med. 2015;7(1):85.

39. Vahdati Nia B, Kang C, Tran MG, Lee D, Murakami S. Meta Analysis of Human AlzGene Database: Benefits and Limitations of Using *C. elegans* for the Study of Alzheimer’s Disease and Co-morbid Conditions. Front Genet. 2017 8:55. doi: 10.3389/fgene.2017.00055.

40. Le D, Brown L, Malik K, Murakami S. Two Opposing Functions of Angiotensin-Converting Enzyme (ACE) That Links Hypertension, Dementia, and Aging. Int J Mol Sci. 2021 22(24):13178. doi: 10.3390/ijms222413178.

41. López-Otín C, Blasco MA, Partridge L, Serrano M, Kroemer G. Hallmarks of aging: An expanding universe. Cell. 2023 186(2):243–278. doi: 10.1016/j.cell.2022.11.001.

42. D’Angelo, M. A., & Hetzer, M. W. (2008). Structure, dynamics and function of nuclear pore complexes. Trends in Cell Biology, 18(10), 456–466. 10.1016/j.tcb.2008.07.009

43. Martins F, Sousa J, Pereira CD, da Cruz E, Silva OAB, Rebelo S. Nuclear envelope dysfunction and its contribution to the aging process. Aging Cell. 2020 19(5):e13143. doi: 10.1111/acel.13143.

44. Eriksson M, Brown WT, Gordon LB, Glynn MW, Singer J, Scott L, Erdos MR, Robbins CM, Moses TY, Berglund P, Dutra A, Pak E, Durkin S, Csoka AB, Boehnke M, Glover TW, Collins FS. Recurrent de novo point mutations in lamin A cause Hutchinson-Gilford progeria syndrome. Nature. 2003 423(6937):293–8. doi: 10.1038/nature01629.

45. Cenni V, Capanni C, Mattioli E, Schena E, Squarzoni S, Bacalini MG, Garagnani P, Salvioli S, Franceschi C, Lattanzi G. Lamin A involvement in ageing processes. Ageing Res Rev. 2020 62:101073. doi: 10.1016/j.arr.2020.101073.

46. Friedrich K, Lee L, Leistritz DF, Nürnberg G, Saha B, Hisama FM, Eyman DK, Lessel D, Nürnberg P, Li C, Garcia-F-Villalta MJ, Kets CM, Schmidtke J, Cruz VT, Van den Akker PC, Boak J, Peter D, Compoginis G, Cefle K, Ozturk S, López N, Wessel T, Poot M, Ippel PF, Groff-Kellermann B, Hoehn H, Martin GM, Kubisch C, Oshima J. WRN mutations in Werner syndrome patients: genomic rearrangements, unusual intronic mutations and ethnic-specific alterations. Hum Genet. 2010 128(1):103–11. doi: 10.1007/s00439-010-0832-5.

47. Martin GM, Hisama FM, Oshima J. Review of How Genetic Research on Segmental Progeroid Syndromes Has Documented Genomic Instability as a Hallmark of Aging But Let Us Now Pursue Antigeroid Syndromes! J Gerontol A Biol Sci Med Sci. 2021 Jan 18;76(2):253–259. doi: 10.1093/gerona/glaa273. PMID: 33295962; PMCID: PMC7812512.

